# Daughter centrioles assemble preferentially towards the nuclear envelope in *Drosophila* syncytial embryos

**DOI:** 10.1101/2021.11.17.468935

**Authors:** Neil H. J. Cunningham, Imène B. Bouhlel, Paul T. Conduit

**Affiliations:** Department of Zoology, University of Cambridge, Downing Street, Cambridge, CB2 3EJ; Université de Paris, CNRS, Institut Jacques Monod, F-75006, Paris, France

**Keywords:** Centrosome, centriole, centriole duplication, Drosophila, microtubules

## Abstract

Centrosomes are important organisers of microtubules within animal cells. They comprise a pair of centrioles surrounded by the pericentriolar material (PCM), which nucleates and organises the microtubules. To maintain centrosome numbers, centrioles must duplicate once and only once per cell cycle. During S-phase, a single new “daughter” centriole is built orthogonally on one side of each radially symmetric “mother” centriole. Mis-regulation of duplication can result in the simultaneous formation of multiple daughter centrioles around a single mother centriole, leading to centrosome amplification, a hallmark of cancer. It remains unclear how a single duplication site is established. It also remains unknown whether this site is pre-defined or randomly positioned around the mother centriole. Here, we show that within *Drosophila* syncytial embryos daughter centrioles preferentially assemble on the side of the mother facing the nuclear envelope, to which the centrosomes are closely attached. This positional preference is established early during duplication and remains stable throughout daughter centriole assembly, but is lost in centrosomes forced to lose their connection to the nuclear envelope. This shows that non-centrosomal cues influence centriole duplication and raises the possibility that these external cues could help establish a single duplication site.

## Introduction

Centrosomes are important microtubule organising centres (MTOCs) within animal cells, best known for organising the mitotic spindle poles during cell division (Conduit et al., 2015b). They typically comprise an older “mother” and younger “daughter” pair of barrel-shaped microtubule-based centrioles. While centriole structure varies between species and cell type (Loncarek and Bettencourt-Dias, 2018), they all display a 9-fold radial symmetry, with an inner “cartwheel” structure supporting the assembly of 9 microtubule triplets, doublets or singlets that make up the centriole wall. The mother centriole recruits and organises a surrounding pericentriolar material (PCM), which contains the necessary microtubule-associating and signalling proteins required for centrosome function (Woodruff et al., 2014). The mother centriole also templates the assembly of the daughter centriole in a process called centriole duplication (Banterle and Gönczy, 2017; Firat-Karalar and Stearns, 2014; Fu et al., 2015). This occurs after cell division, when each daughter inherits a single centrosome containing a disengaged mother-daughter centriole pair. The daughter centriole is converted into a mother and both mothers support the orthogonal assembly of a new daughter centriole at their proximal end during S-phase. The two mother-daughter centriole pairs break apart during G2/M-phase to form two centrosomes, which mature by recruiting more PCM in preparation for mitosis. During mitosis, the two centrosomes each organise one pole of the bipolar spindle and towards the end of mitosis the centrioles disengage in preparation for a new round of duplication in the following cell cycle.

In most cell types, centrioles duplicate once per cell cycle during S-phase and it is this “once and only once” duplication event that maintains centrosome numbers through multiple cell divisions (Nigg and Holland, 2018). Failure to duplicate the centrioles during S-phase results in the inherence of a centrosome with a single centriole, which cannot then split to form two centrosomes. This leads to monopolar spindle formation and cell cycle arrest. In contrast, multiple centrosomes form if mother centrioles template the assembly of more than one daughter centriole and this leads to multipolar spindle formation in the next cell cycle. Multipolar spindles can result in cell death or they can be transformed into bipolar spindles that harbour erroneous kinetochore attachments, leading to lagging chromosomes and chromosome instability (Basto et al., 2008; Ganem et al., 2009; Godinho and Pellman, 2014; Nigg and Holland, 2018). Centrosome amplification is strongly associated with cancer progression, with chromosome instability and increased centrosome signalling being possible causal links (Anderhub et al., 2012; Basto et al., 2008; Denu et al., 2016; Godinho et al., 2014; Godinho and Pellman, 2014; Mittal et al., 2021; Salisbury et al., 2004).

Seminal studies in *C. elegans* identified a core set of proteins necessary for centriole duplication: the kinase ZYG-1 and the large coiled-coil proteins SPD-2, SAS-4, SAS-5 and SAS-6 (Delattre et al., 2006; Leidel and Gönczy, 2003; O’Connell et al., 2001; Pelletier et al., 2006). Homologues in *Drosophila* (Sak/Plk4, Spd-2, Sas-4, Ana2 and Sas-6) and human cells (PLK4, CEP192, CPAP, STIL and SAS-6) were subsequently identified and, with the exception of *Drosophila* Spd-2 (Dix and Raff, 2007), shown to also be essential for centriole duplication (Basto et al., 2006; Bettencourt-Dias et al., 2005; Dammermann et al., 2004; Habedanck et al., 2005; Kim et al., 2013; Leidel et al., 2005; Sonnen et al., 2013; Stevens et al., 2010; Tang et al., 2011; Terra et al., 2005). The role of worm SPD-2, which is to recruit ZYG1/PLK4, is played instead by *Drosophila* Asterless (Asl) (Blachon et al., 2008; Dzhindzhev et al., 2010), and the human homologues of Asl (CEP152) is also required for centriole duplication (Blachon et al., 2008; Cizmecioglu et al., 2010; Hatch et al., 2010), functioning together with the human homologue of SPD-2 (CEP192) to recruit PLK-4 (Kim et al., 2013; Park et al., 2014; Sonnen et al., 2013).

A large number of studies are producing a clear picture about how each of these proteins contributes to centriole assembly (reviewed in (Arquint and Nigg, 2016; Firat-Karalar and Stearns, 2014; Fu et al., 2015; Yamamoto and Kitagawa, 2021). In essence, CEP192/SPD-2 and/or CEP152/Asl recruit the master kinase PLK-4 to the wall of the mother centriole where it regulates the recruitment of STIL/Ana2 and SAS-6 and then CPAP/Sas-4 to form the daughter centriole. A key feature is that daughter centriole assembly occurs on only one side of the radially symmetric mother centriole, and this relies on localising PLK4, SAS-6 and STIL/Ana2 to a single spot on the side of the mother. The problem is that CEP192/SPD-2 and CEP152/Asl localise as a ring around the mother centriole and thus PLK4 is also initially recruited in a ring-like pattern (Kim et al., 2013; Park et al., 2014; Sonnen et al., 2013). In order for just a single daughter centriole to form, this ring of PLK4 must therefore be converted to a ‘dot’, which marks the site of centriole duplication. Failure of PLK4 to undergo this ‘ring-to-dot’ conversion results in multiple daughter centrioles forming around the mother centriole and this leads to centrosome amplification (Brownlee et al., 2011; Habedanck et al., 2005; Klebba et al., 2013; Kleylein-Sohn et al., 2007; Ohta et al., 2014). Ring-to-dot conversion of PLK4 is thought to be largely self-controlled, as it involves the auto-phosphorylation of a degron within PLK4 (Cunha-Ferreira et al., 2013, 2009; Guderian et al., 2010; Holland et al., 2010; Klebba et al., 2013; Rogers et al., 2009; Sillibourne et al., 2010), and could also depend on the ability of PLK4 to self-assemble, a property that is regulated by auto-phosphorylation and that protects PLK4 from degradation (Gouveia et al., 2018; Park et al., 2019; Yamamoto and Kitagawa, 2019). Nevertheless, ring-to-dot conversion is likely also influenced by the binding of STIL/Ana-2, which increases PLK4 activity (Arquint et al., 2015; Moyer et al., 2015) and protects PLK4 from degradation (Arquint et al., 2015; Ohta et al., 2014). In human cells, PLK4 is observed as an asymmetric punctate ring prior to the recruitment of STIL, suggesting that initial symmetry breaking is independent of STIL, although the full ring-to-dot conversion occurs only once STIL and SAS-6 have been recruited (Kim et al., 2013; Ohta et al., 2018, 2014; Park et al., 2014; Yamamoto and Kitagawa, 2019). In flies, Ana2 recruitment is the first observed symmetry breaking event (Dzhindzhev et al., 2017). Mathematical models can explain how the properties of PLK4, with or without the help of STIL/Ana2, can lead to the symmetry breaking ring-to-dot transition (Leda et al., 2018; Takao et al., 2019).

While various studies have focussed on understanding how symmetry breaking is achieved, it remains unknown whether the site of daughter centriole assembly is randomly assigned or not. We decided to investigate this using *Drosophila* syncytial embryos as a model system. These embryos go through rapid and near-synchronous rounds of S-phase and then mitosis with no intervening gap phases. The nuclear envelope does not fully break down during mitosis and the centrosomes remain closely attached to the nuclear envelope throughout each cycle. At the end of mitosis / start of S-phase, mother and daughter centrioles separate with the daughter converting to a mother and both centrioles quickly migrate around the nuclear envelope to form two new centrosomes that will organise the next round of mitosis. During S-phase, each mother centriole templates the formation of a new daughter centriole, with only the mother centriole organising PCM (Conduit et al., 2015a, 2010). Towards the end of mitosis, the centrioles disengage and the daughter centrioles are converted to mothers by the addition of Asl, allowing them to begin recruiting PCM and initiate centriole duplication in the next cycle (Conduit et al., 2010; Novak et al., 2014).

Using a dual-colour FRAP approach along with super-resolution Airyscan imaging, we show here that daughter centrioles preferentially assemble on the side of the mother centriole facing the nuclear envelope. By tracking duplication events throughout S-phase, we show that this preferential positioning of the daughter centriole with respect to the nucleus occurs from the early stages of centriole formation and remains relatively stable throughout the cycle. Using a point mutation in the key PCM protein Centrosomin (Cnn), we show that this preferential positioning towards the nuclear envelope is lost in centrosomes that have detached from the nuclear envelope. Collectively, these observations suggest that the site of centriole duplication is influenced by the nuclear envelope and raise the possibility that cues external to the centriole duplication machinery may influence and help control centriole duplication.

## Results

### The site of daughter centriole assembly is non-random with respect to cell geometry

To address whether the site of daughter centriole formation is pre-defined or randomly assigned during centriole duplication, we turned to the *Drosophila* syncytial embryo. In these embryos hundreds of nuclei and centrosomes undergo rapid cycles of division (∼8-15 min per cycle) in near synchrony, alternating between S-phase and M-phase without gap phases. At around division cycle 9 the nuclei and centrosomes migrate to the cell cortex and their divisions can be readily imaged with a fluorescence-based microscope until they pause in cycle 14. Mitotic spindles form parallel to the cortex such that they align along the X-Y imaging plane. The mother centrioles also have a regular alignment; their proximal-distal (end-to-end) axis is aligned orthogonally to the spindle axis such that mother centrioles point along the Z imaging axis. Newly forming daughter centrioles grow along the X-Y imaging axis. This regular alignment of the centrioles in theory allows one to record the position of the daughter centriole relative to other cellular structures, such as the mitotic spindle axis. *Drosophila* centrioles are relatively small, however, meaning that duplicating mother-daughter centriole pairs cannot be resolved using “standard” confocal microscopy. We therefore developed a method to estimate the location of the centrioles within an engaged mother-daughter centriole pair by performing dual-colour Fluorescence Recovery After Photobleaching (FRAP) experiments. This relies on the fact that PCM proteins, such as Spd-2, Asl or Cnn, are dynamically recruited around the mother, but not the daughter, centriole, while the centriole protein Sas-4 is dynamically recruited to the growing daughter, but not the mother, centriole (Conduit et al., 2015a). By tagging a PCM protein and Sas-4 with different coloured fluorophores and then photobleaching during S-phase, the centroids of the recovering fluorescent signals can be used to estimate the relative positions of the mother (PCM signal) and daughter centrioles (Sas-4 signal) (Figure 1A). We used this approach to compare the position of the growing daughter centriole relative to the mother centriole and the future spindle axis (Figure 1B).

**Figure 1.**
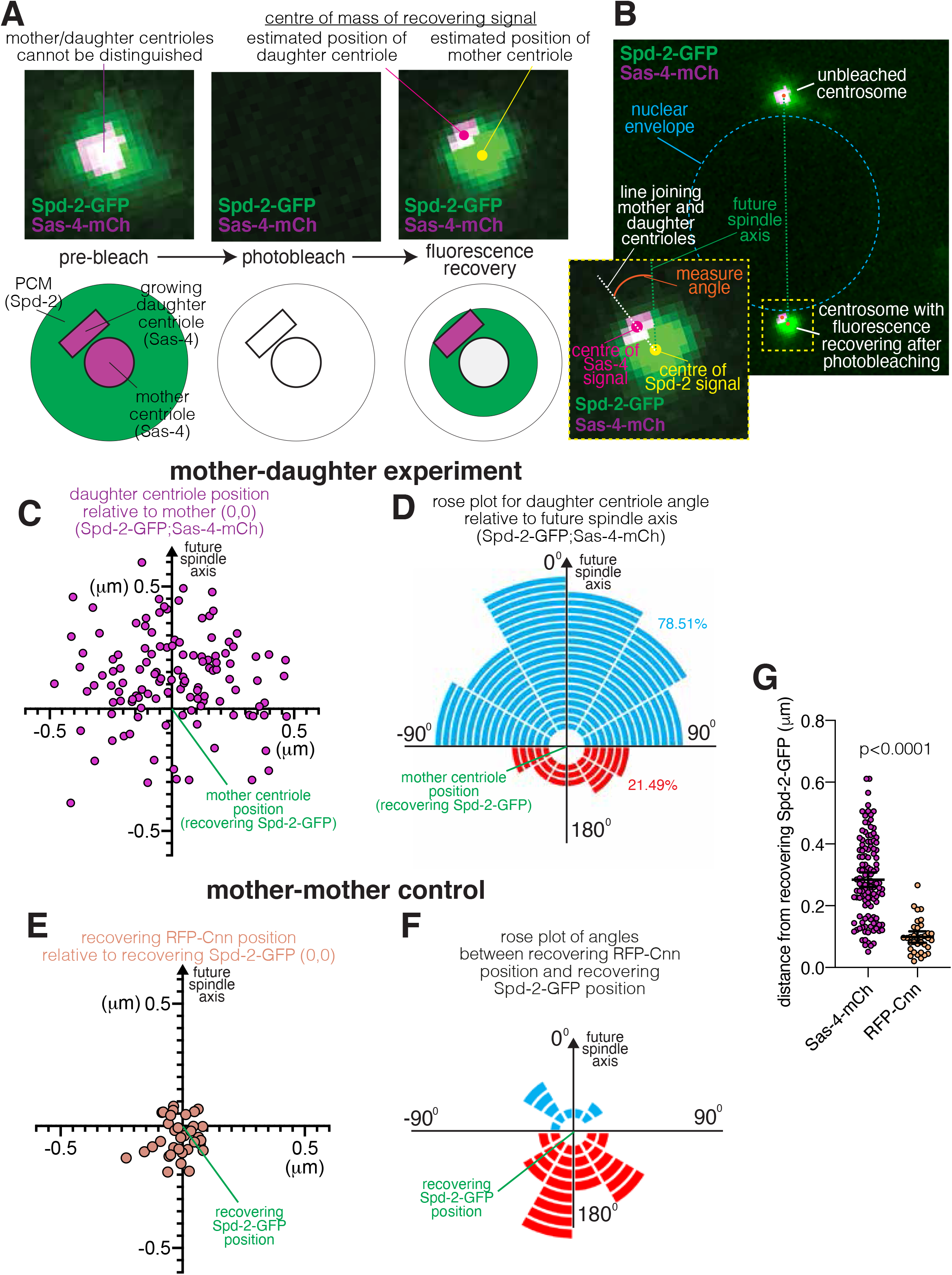
Analysis of dual-colour FRAP data reveals that the site of daughter centriole assembly is non-random. (**A**) Confocal images show a centrosome within an embryo expressing Spd-2-GFP (green) and Sas-4-mCherry (magenta) prior to photobleaching (left), immediately after photobleaching (centre), and after fluorescence recovery (right). The diagrams below are cartoon representations of how the proteins behave before and after photobleaching. Note that the recovering Sas-4-mCherry signal (daughter centriole) is offset from the centre of the recovering Spd-2-GFP signal (mother centriole). (**B**) Confocal image shows a pair of centrosomes (top unbleached, bottom recovering from bleaching) on opposite sides of the nuclear envelope (mid-late S-phase). The nuclear envelope and how angles from the future spindle axis are calculated are indicated. (**C**) Graph displays the estimated positions of daughter centrioles (magenta circles) relative to the estimated position of their respective mother centrioles (position 0,0 on the graph) and the future spindle axis (positive y-axis) obtained from Spd-2-GFP (mother) Sas-4-mCherry (daughter) data. (**D**) Rose plot representing the angle at which daughter centrioles (marked by Sas-4-mCherry) form in relation to the future spindle axis (0°). Each segment corresponds to a single duplication event. Blue and red segments indicate daughter centriole assembly occurring less than or more than 90° from the future spindle axis, respectively. (**E**) Graph displays the positions of the centre of recovering RFP-Cnn signal relative to recovering Spd-2-GFP signal (position 0,0 on the graph) and the future spindle axis (positive y-axis) obtained from the control Spd-2-GFP (mother) RFP-Cnn (mother) data. (**F**) Rose plot (as in (D)) representing the angle relative to the future spindle axis (0°) formed by a line running between the recovering Spd-2-GFP and RFP-Cnn signals. (**G**) Graph showing the distance between the centre of the recovering Spd-2-GFP signal (mother centriole) and the recovering Sas-4-mCherry signal (daughter centriole, magenta) or the recovering RFP-Cnn signal (mother centriole). The datasets were compared using a Mann-Whitney test.

To begin with, we used Spd-2-GFP and Sas-4-mCherry as our mother and daughter centriole markers, respectively. We photobleached either one centrosome from a separating centrosome pair during early S-phase (when Sas-4 starts to be incorporated at the newly forming daughter centriole) or we photobleached a single centrosome in late M-phase, just prior to centrosome splitting, daughter centriole assembly and Sas-4 recruitment, and monitored the two resulting centrosomes in the following S-phase. Both cases result in centrosomes where Spd-2-GFP recovers only around the mother centriole and Sas-4-mCherry recovers only at the growing daughter centrioles during S-phase, but the latter case generates two centrosomes that can be analysed. We recorded the centroids of the recovering fluorescent signals in mid to late S-phase once the centrosomes had reached their final positions on the opposite side of the nuclear envelope. Waiting until the centrosomes had fully separated allowed us to use the future spindle axis (a line drawn between the paired centrosomes) as a spatial reference point with which to compare the position of daughter centriole assembly (Figure 1B). We analysed a total of 121 centrosomes from 16 embryos and collated the results. Strikingly, the positions of daughter centrioles were not evenly distributed relative to the future spindle axis (positive Y axis in Figure 1C). A frequency distribution of the angles of the daughter centrioles relative to the future spindle axis showed displayed a Normal distribution around the 0° angle (Figure S1A,B) (passed all 4 normality tests in Prism) i.e. the daughter centrioles had a preference to be close to the 0° angle and were not evenly distributed around the mother centriole (Chi-square=44.52, df=11, p<0.0001), as would be expected if daughter centriole positioning were random. The data can also be represented by a Rose Plot, where each segment corresponds to a duplication event and its position corresponds to the angle from the future spindle axis (Figure 1D). 95 of 121 (78.51%) daughter centrioles were assembled within 90 degrees of the future spindle axis (blue segments, Figure 1D), while only 26 (21.49%) were assembled more than 90 degrees from the future spindle axis (red segments, Figure 1F) (Binomial Wilson/Brown test, p<0.0001). The distribution of daughter centriole positions was not due to microscope induced misalignment of the green and red channels: auto-fluorescent beads were used to correct for microscope-induced offset between the channels (as in (Conduit et al., 2015a)); and the data was taken from multiple nuclei/centrosome pairs, all of which have different orientations with respect to the X-Y axes of the microscope. Moreover, we observed a more random and non-Normal distribution of angles when imaging the fluorescence recovery of two PCM proteins, Spd-2-GFP and RFP-Cnn, which are expected to be closely aligned (Figure 1E,F; Figure S1C,D). Indeed, the positions of the recovering RFP-Cnn signals relative to the recovering Spd-2-GFP signals were much closer together with the mean distance between these signals (0.099μm) being significantly shorter than the mean distance between the recovering Spd-2-GFP (mother) and Sas-4-GFP (daughter) signals (0.284μm) (Figure 1G). We also repeated the experiment using a green version of Sas-4 (Sas-4-GFP) and a different mother centriole marker (Asl-mCherry) on a different microscope and again found that the positions of daughter centriole assembly were not evenly distributed relative to the future spindle axis (Figure S1E), that the angles from the future spindle axis were Normally distributed around 0° (Figure S1F,G), that a much higher proportion of daughter centrioles assembled within 90 degrees of the future spindle axis (Figure S1H), and that the distance between the recovering signals was similar to that for the Spd-2-GFP/Sas-4-mCherry data (Figure S1I). Collectively, this data shows that the positioning of daughter centriole assembly in *Drosophila* syncytial embryos is non-random with respect to cellular geometry.

**Figure S1.**
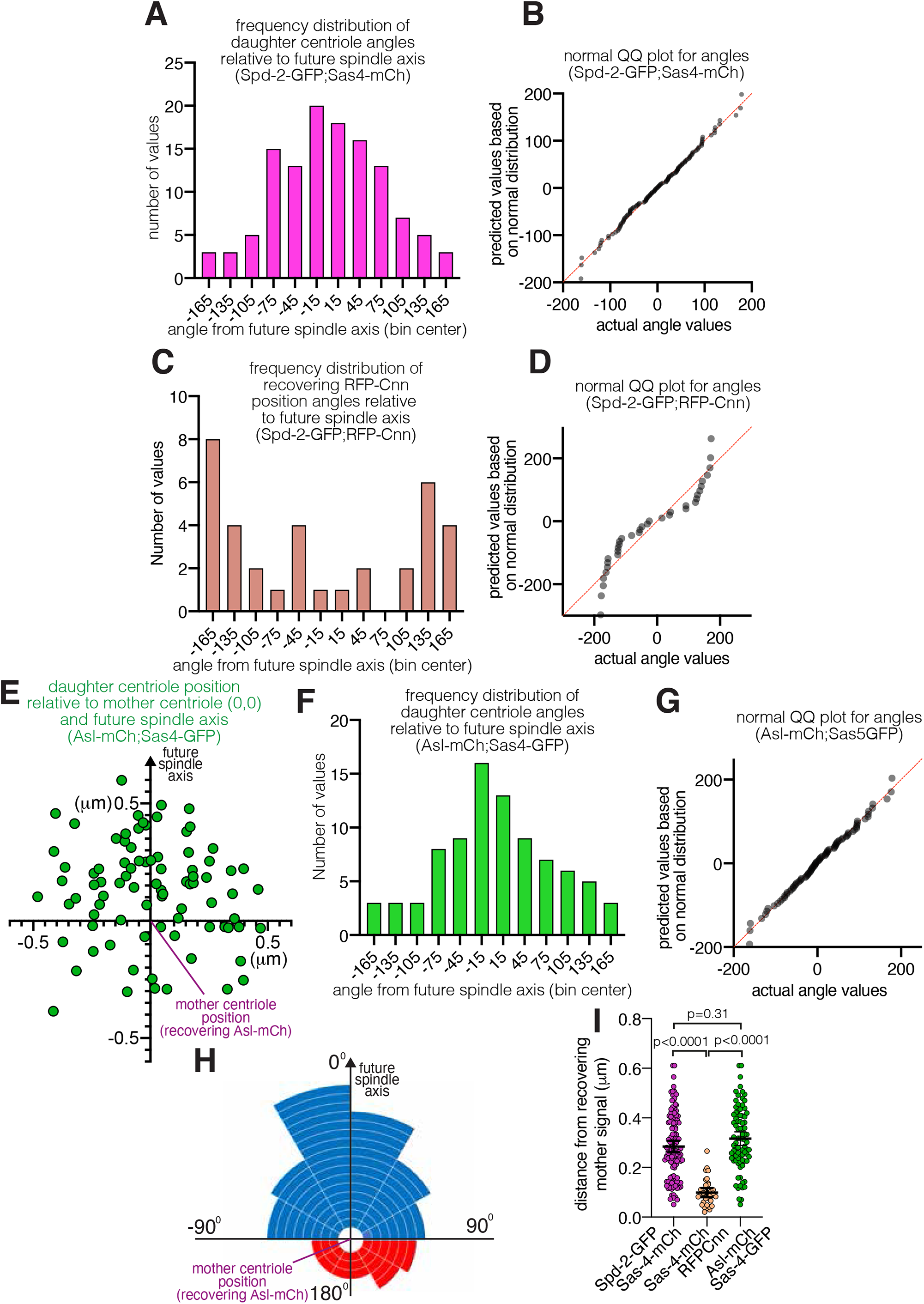
Further analysis of dual-colour FRAP data supports the finding that the site of daughter centriole assembly is non-random. (**A**) Frequency distribution of the angles at which daughter centrioles (marked by Sas-4-mCherry) form in relation to the future spindle axis (0°). (**B**) Normal QQ plot showing that the angles in (A) conform well to a normal distribution. (**C**) Frequency distribution of the angles at which the recovering RFP-Cnn fluorescence is positioned in relation to the future spindle axis (0°). (**D**) Normal QQ plot showing that the angles in (B) do not conform well to a normal distribution. (**E**) Graph displays the estimated positions of daughter centrioles (green circles) relative to the estimated position of their respective mother centrioles (position 0,0 on the graph) and the future spindle axis (positive y-axis) obtained from Asl-mCherry (mother) Sas-4-GFP (daughter) data. (**F**) Frequency distribution of the angles at which daughter centrioles (marked by Sas-4-GFP) form in relation to the future spindle axis (0°). (**G**) Normal QQ plot showing that the angles in (F) conform well to a normal distribution. (**H**) Rose plot representing the angle at which daughter centrioles (marked by Sas-4-GFP) form in relation to the future spindle axis (0°). Each segment corresponds to a single duplication event. Blue and red segments indicate daughter centriole assembly occurring less than or more than 90° from the future spindle axis, respectively. (**I**) Graph showing the distance between the estimated positions of mother and daughter centrioles (left and right datasets) or two different estimations of the mother centriole (central dataset) in the different imaging conditions used, as indicated. Note that data for the datasets on the left and in the centre have been re-plotted from Figure 1G to allow comparison to the dataset on the right. Datasets were compared to each other using a one-way ANOVA Kruskal-Wallis test.

### The non-random position of daughter centriole assembly is dependent on centrosome association with the nuclear envelope

In *Drosophila* syncytial embryos, the centrosomes are tightly associated with the nuclear envelope via nuclear envelope associated Dynein (Robinson et al., 1999). Thus, the observation that daughter centrioles form preferentially within 90° of the future spindle axis also meant that they were preferentially positioned on the side of the mother centriole facing the nuclear envelope. This raised the intriguing possibility that the nuclear envelope might influence the position of daughter centriole assembly. To test this, we wanted to examine the position of daughter centriole assembly in centrosomes that had detached from the nuclear envelope. We knew that Threonine 1133 within the PCM protein Cnn is important for Cnn to oligomerise and form a PCM scaffold (Feng et al., 2017) and our unpublished observations had shown that substituting Threonine 1133 with Alanine partially perturbs scaffold formation and the ability of centrosomes to remain attached to the nuclear envelope (see also Figure 2A). We therefore generated a stock co-expressing Sas-4-mCherry and a GFP-Cnn-T1133A to analyse daughter centriole position in attached versus detached centrosomes. The detached centrosomes in Cnn-T1133A mutants normally remain relatively close to the nuclear envelope, do not fall into the embryo centre, and form a spindle pole in during the following mitosis. Nevertheless, they often do not fully migrate around the nucleus (Figure 2A). Thus, instead of using the line between paired centrosomes as a reference point for the angle of daughter centriole assembly, we used a line drawn between the mother centriole and the centre of the nucleus (visualised due to the exclusion of fluorescence molecules), which we hereafter refer to as the nuclear axis (Figure 2A,B).

**Figure 2.**
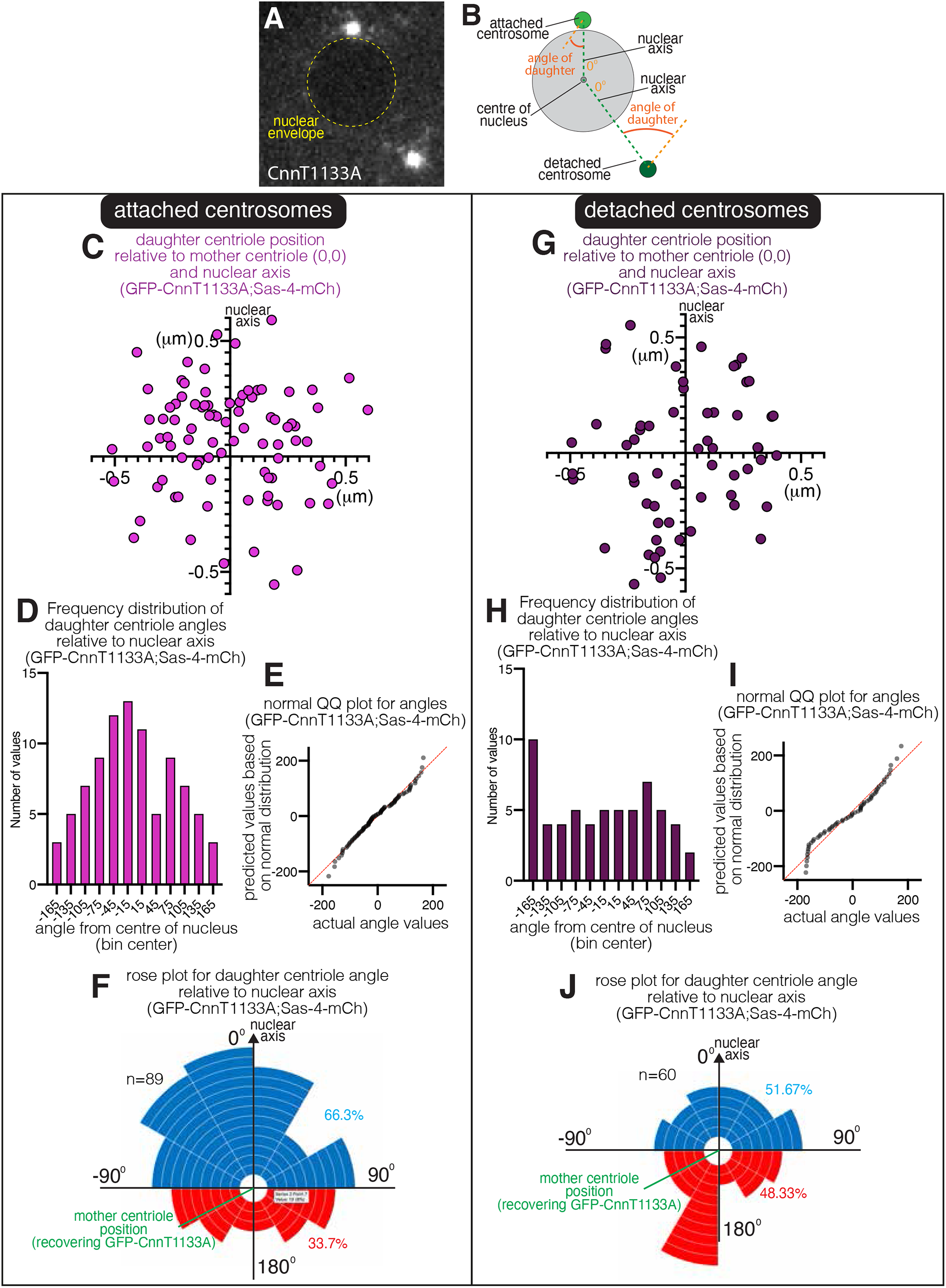
The site of daughter centriole assembly is random in centrosomes that have detached from the nuclear envelope. (**A**,**B**) Confocal image (A) and cartoon representation (B) show a pair of centrosomes in S-phase within an embryo expressing GFP-Cnn-T1133A (grayscale). Note that one centrosome is attached to and one centrosome is detached from the nuclear envelope. Cartoon in (B) indicates how the angles of daughter centriole assembly from the nuclear axis were measured. (**C-J**) Graphs display results from analysing the estimated position of daughter centrioles relative to the estimated position of their respective mother centrioles (position 0,0 on the graph) and the nuclear axis (positive y-axis) in centrosomes that have either remained attached to (C-F) or that have detached from (D-J) the nuclear envelope within embryos expressing GFP-Cnn-T1133A and Sas-4-mCherry. Estimated positions of the daughter centrioles were determined from analysing the centre of fluorescence recovery of GFP-Cnn-T1133A (mother) and Sas-4-mCherry (daughter). Graphs in (C) and (G) show the estimated positions of the daughter centrioles; (D) and (H) are frequency distributions of the angles at which daughter centriole form in relation to the nuclear axis (0°); (E) and (I) are normal QQ plot showing that the angles in (E), but not in (I), conform well to a Normal distribution; Rose plots in (F) and (J) represent the angle at which daughter centrioles form in relation to the mother centriole and the nuclear axis (0°). Each segment corresponds to a single duplication event. Blue and red segments indicate daughter centriole assembly occurring less than or more than 90° from the nuclear axis, respectively.

We photobleached centrosomes in late mitosis and monitored the fluorescence recovery during the following S-phase, noting which centrosomes had separated from the nuclear envelope and which had not. Importantly, the daughter centrioles within centrosomes that had remained attached to the nuclear envelope still displayed a preference to assemble on the side of the mother facing the nuclear envelope (Figure 2C-F), showing that perturbation of the PCM via Cnn’s T1133A mutation did not indirectly affect daughter centriole positioning. In these attached centrosomes, the estimated position of the daughter centrioles displayed a similar non-even distribution to that observed in the analyses above for Spd-2-GFP;Sas-4-mCherry and Asl-mCherry; Sas-4-GFP (compare Figures 1C, 2C and Figure S1E). The measured angles of daughter centriole formation were normally distributed around 0° (Figure 2D,E) (passed all 4 Normality tests in Prism) and a Rose Plot graph highlighted how 66.3% (59 of 89) daughter centrioles were positioned within 90 degrees of 0° (Figure 2F) (Binomial Wilson/Brown test, p<0.01). In contrast to the attached centrosomes, the daughter centrioles within centrosomes detached from the nuclear envelope did not display a preference to assemble on the side of the mother facing the nuclear envelope (Figure 2G-J). The estimated position of these daughter centrioles was more evenly spread around the mother centriole (Figure 2G) and the angles at which they assembled relative to the nuclear axis were not normally distributed around 0° (Figure 2H,I) (Failed 3 of 4 Normality tests in Prism) and were not significantly different from a random distribution (Chi-square=8.4, df=11, p=0.68). Moreover, there was no preference for the centrioles to form within 90 degrees of the nuclear axis, with similar numbers of daughter centrioles forming within 90 degrees (31/60) and more than 90 degrees (29/60) from the nuclear axis (Figure 2J) (Binomial Wilson/Brown test, p=0.90).

It was possible that the perceived loss of preference for the daughter centriole to form towards the nuclear axis in detached Cnn-T1133A centrosomes could have been an indirect effect of defects in centriole orientation with respect to the imaging axis i.e. detached centrosomes may tilt such that their daughter centrioles do not grow along the X-Y imaging axis, causing increased noise and a possible randomising effect in the data. We ruled this out in two different ways. First, we compared the frequency at which GFP-Cnn-T1133A displayed a “central hole” at attached and detached centrosomes. Cnn molecules surround the mother centriole such that, with sufficient X-Y spatial resolution, a “hole” in the centre of the Cnn fluorescence signal can be observed (e.g. top panels in Figure 3A, B). We reasoned that this central hole would be observed only in centrosomes that had their mother centriole pointing normally along the Z imaging axis. We imaged fixed embryos in S-phase expressing GFP-Cnn-T1133A and Asl-mCherry (which labels only mother centrioles during S-phase) on a Zeiss Airyscan 2 microscope, which increases X-Y spatial resolution to up to 120nm, and quantified the frequency of “clear”, “partial”, or “no clear” central holes in attached (Figure 3A) versus detached (Figure 3B) centrosomes. Out of a total of 112 centrosomes from 3 embryos, 83 were attached and 29 were detached. Of the 83 attached centrosomes, 38 (45.8%) displayed a clear central hole, 25 (30.1%) displayed a partial central hole, and 20 (24.1%) displayed no clear central hole (Figure 3C). These percentages were similar in detached centrosomes. Of the 29 detached centrosomes, 12 (41.4%) displayed a clear doughnut-like pattern, 9 (31.0%) displayed a partial doughnut-like pattern, and 8 (27.6%) displayed no clear doughnut-like pattern (Figure 3C). There was no significant difference between the categorisation of these attached and detached centrosomes (Chi-square = 0.204, df=2, p=0.903), suggesting that detached centrosomes are not mis-oriented compared to attached centrosomes. To further support this finding, we used the previous Spd-2-GFP/Sas-4-mCherry FRAP data (Figure 2C,G) to compare the median estimated distances between mother and daughter centrioles in attached (0.30μm) versus detached (0.33μm) centrosomes and found there was no significant difference (Figure 3D; Mann-Whitney, p=0.26). The distance would in theory be shorter in detached centrosomes if they were misoriented. Thus, the data suggests that mother centrioles within centrosomes that have detached from the nuclear envelope remain aligned along the Z imaging axis. We therefore conclude that, unlike in attached centrosomes, daughter centrioles within detached centrosomes do not form preferentially towards the nuclear envelope and that the nucleus somehow influences daughter centriole positioning.

**Figure 3.**
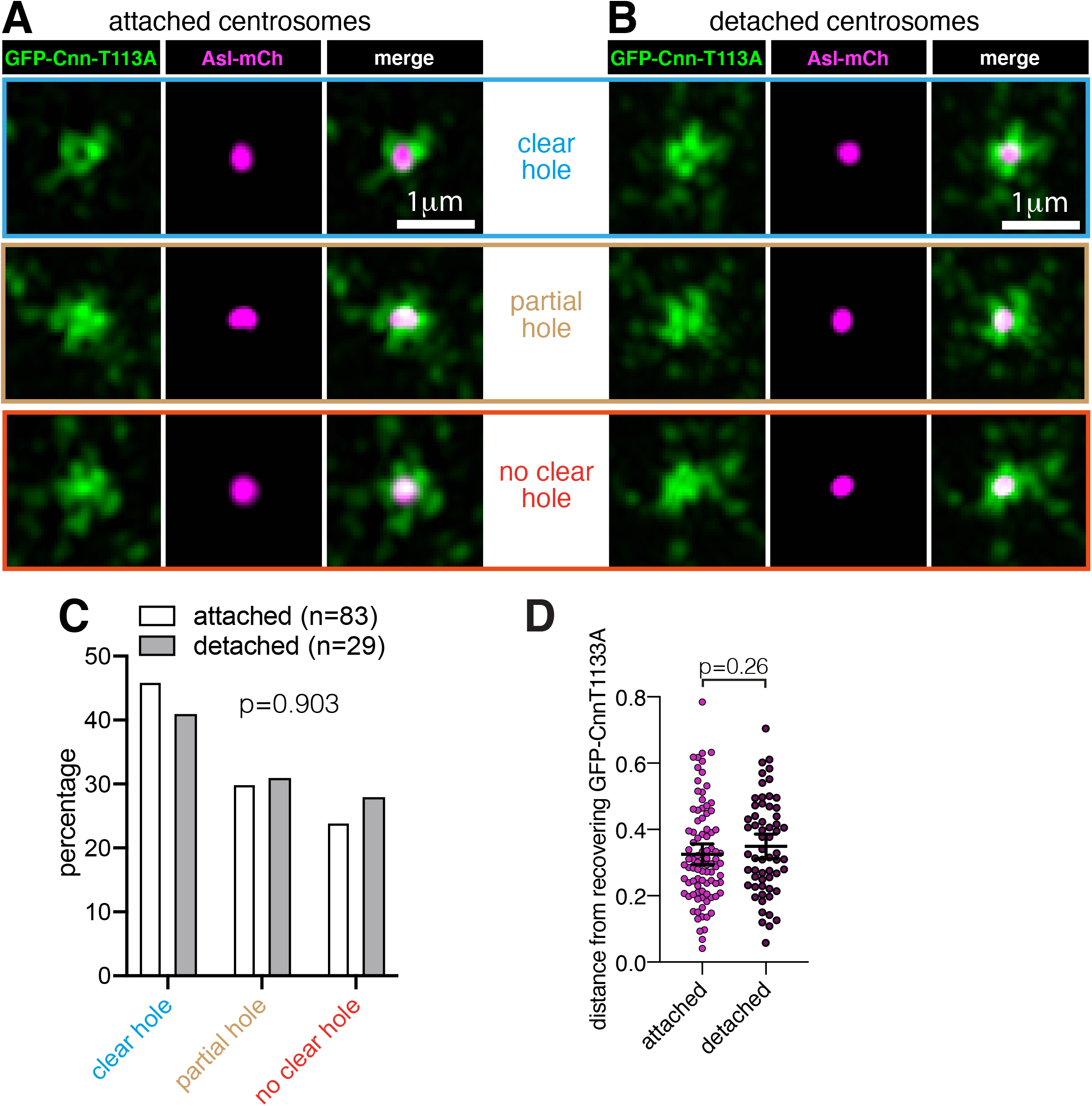
Cnn-T1133A centrosomes that have detached from the nuclear envelope remain correctly oriented with respect to the imaging axis. (**A**,**B**) Airyscan images of centrosomes that are either attached to (A) or detached from (B) the nuclear envelope within embryos expressing GFP-Cnn-T1133A and Sas-4-mCherry in a *cnn* null mutant background. Examples with a clear central hole (top panels), a partial central hole (middle panels), and a no clear central hole (bottom panels) are shown. (**C**) Graph shows the percentage of each centrosome type in either attached or detached centrosomes, as indicated. Datasets were compared using a Chi-squared contingency analysis. (**D**) Graph shows the distances between the estimated positions of mother and daughter centrioles from the Spd-2-GFP/Sas-4-mCherry FRAP data in either attached or detached centrosomes, as indicated. The datasets were compared using a Mann-Whitney test.

### The positioning of daughter centriole assembly is consistent through time

To estimate the position of daughter centrioles from our FRAP data, we had needed to wait until the fluorescent signals had recovered sufficiently in order to take accurate measurements, meaning that we could only assess daughter centriole positioning during mid to late S-phase. We therefore wondered whether the initial steps of daughter centriole formation occur with a positional preference, or whether they occur in a random position with the daughter centriole rotating to face the nuclear envelope later in S-phase. To address this, we performed live imaging of duplicating centrosomes throughout S-phase using an Airyscan microscope that enabled us to distinguish two mother and daughter foci of Sas-4-mCherry signal, with the mother centriole localised in the centre of the Spd-2-GFP fluorescence (Figure 4A). Note that the growing daughter centriole rapidly recruits excess Sas-4 (Conduit et al., 2015a) and so appears brighter than the mother for the majority of S-phase, and that Spd-2-GFP, like GFP-Cnn, surrounds the mother centriole and can display a central hole with high enough spatial resolution (certain timepoints in Figure 4A; (Conduit et al., 2014)). Exclusion of cytoplasmic fluorescence can also be used to assess the position of the nuclear envelope (data not shown), which is indicated in blue in Figure 4A (note that centrosomes can migrate over the nucleus, explaining why the paired centrosome in timepoint 1 overlaps the nuclear region).

**Figure 4.**
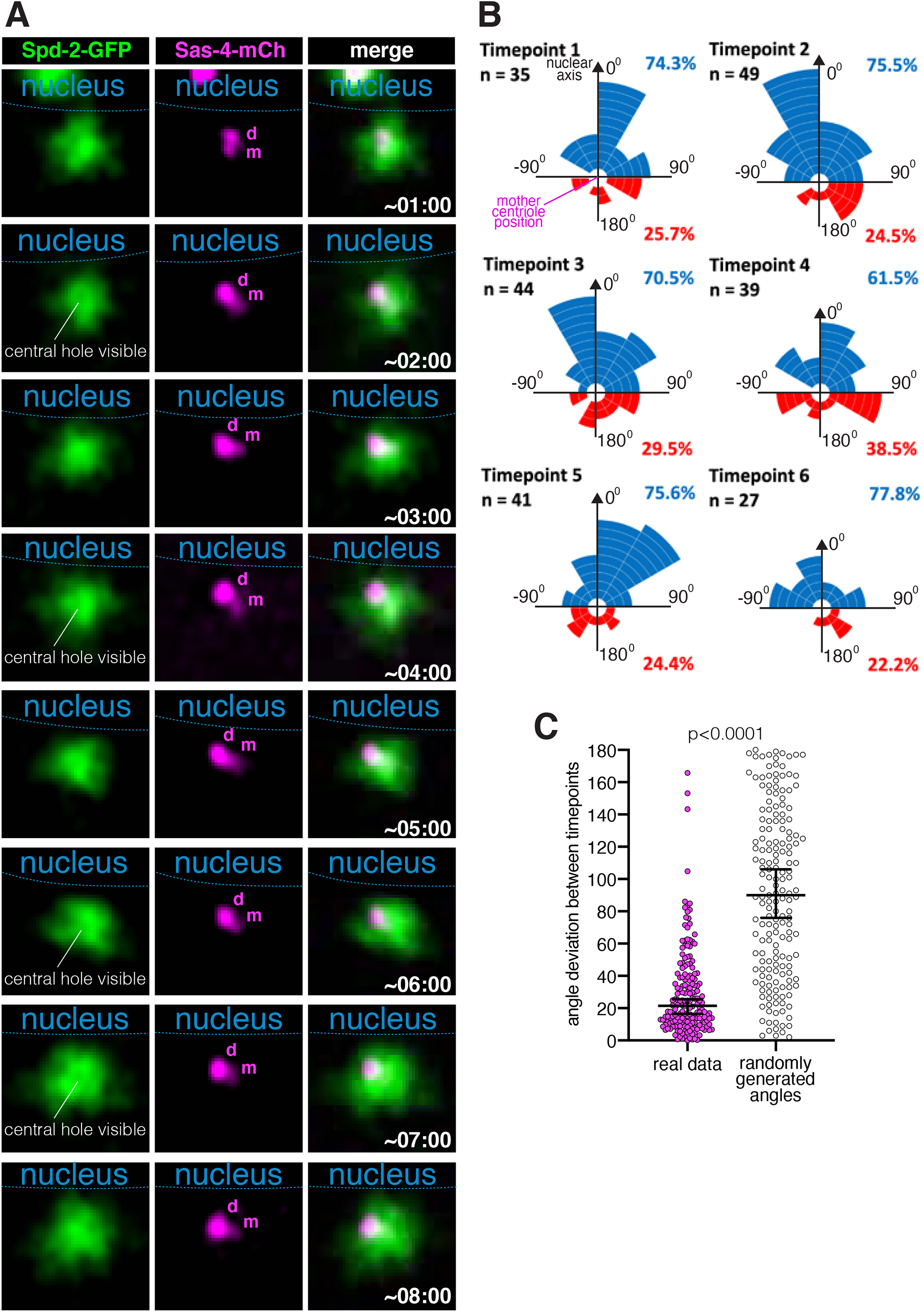
Daughter centrioles initially form preferentially towards the nuclear envelope and retain a stable position throughout S-phase. (**A**) Airyscan images of a centrosome in an embryo expressing Spd-2-GFP (green) and Sas-4-mCherry (magenta) progressing through S-phase. Approximate times after centrosome splitting are indicated – images were collected approximately every minute. The position of the nuclear envelope (as determined by the exclusion of fluorescence from the nucleus) is indicated by the dotted blue line. The Sas-4-mCherry signals for mother (m) and daughter (d) centrioles are also indicated. (**B**) Rose plot graphs display the angle at which daughter centrioles form in relation to the mother centriole and the nuclear axis (0°) as calculated from time-lapse Airyscan images that followed centrosomes throughout S-phase. Each segment corresponds to a single duplication event. Blue and red segments indicate daughter centriole assembly occurring less than or more than 90° from the nuclear axis, respectively. Each rose plot corresponds to a given timepoint, with timepoint 1 occurring ∼1 minute after centrosome splitting and there being a ∼1-minute gap between timepoints. The numbers of events for each timepoint are indicated; this varies due to the varying ability to resolve the two centrioles through time. (**C**) Graph shows the change in the angle of the daughter centriole (angle deviation) with respect to the mother centriole and the nuclear axis that occurred between timepoints from real data (left dataset) or randomly generated angles (right dataset). Each point on the graph represents an individual angle deviation. The median and 95% CIs are shown. The p value indicates that the two datasets have a different distribution (Kolmogorov-Smirnov test).

We followed 72 centrosomes for at least 6 timepoints (∼5 minutes) and collated the data. Note that for most centrosomes, the mother and daughter centrioles within a pair were not resolvable for all 6 timepoints and so the number of measurements per timepoint varied between timepoints. We found that daughter centrioles had a strong preference to assemble on the side of the mother facing the nuclear envelope from the earliest stage of S-phase that the daughter centrioles were visible (timepoint 1, Figure 4B). Moreover, this preference remained throughout the 6 timepoints (Figure 4B). Indeed, we found that daughter centriole positioning relative to the nuclear axis remained quite stable over time. The median angle deviation between timepoints was 21.5°, which is much lower than the median angle deviation expected were the daughter centrioles to be positioned randomly at each timepoint (∼90°). Indeed, the distribution of deviation angles was significantly different from the distribution of random number data (Figure 4C; p<0.0001 Kolmogorov-Smirnov test). Collectively, this data shows that daughter centriole assembly is initiated preferentially on the side of the mother facing the nuclear envelope and that this positioning remains relatively stable throughout daughter centriole assembly.

## Discussion

We have shown that during the mitotic nuclear cycles in *Drosophila* syncytial embryos daughter centrioles preferentially assemble on the side of the mother centriole facing the nuclear envelope. This preferential positioning is lost when centrosomes become detached from the nuclear envelope, raising the intriguing possibility that crosstalk between nuclear-envelope-related factors and the centriole duplication machinery may help to instruct centriole duplication.

A major outstanding question is how PLK4 symmetry breaking is achieved to ensure that only one daughter centriole is formed on the side of the radially symmetric mother centriole (Yamamoto and Kitagawa, 2021). It is known that the PLK4 ring-to-dot transition requires proteasome activity (Ohta et al., 2014), Plk4 activity (Ohta et al., 2018; Park et al., 2019), and phosphorylation of PLK4’s cryptic polo box (Park et al., 2019), suggesting that the auto-catalytic self-destructive properties of PLK4 could regulate the transition (Leda et al., 2018; Park et al., 2019; Takao et al., 2019; Yamamoto and Kitagawa, 2021, 2019). Indeed, computer modelling suggests that PLK4 symmetry breaking can be initiated by the self-organisational properties of PLK4 (Leda et al., 2018; Takao et al., 2019). An initial stochastic break in symmetry could then be enhanced by the binding of STIL (Leda et al., 2018; Takao et al., 2019), which both stimulates PLK4 activity (Moyer et al., 2015) and protects Plk4 from degradation (Arquint et al., 2015; Ohta et al., 2014). The different computer simulations place a difference emphasis on the role of STIL binding (Leda et al., 2018; Takao et al., 2019), but both agree that this is a critical step in completing the ring-to-dot transition. It is intriguing that STIL is able to bind to only a single site on the mother centriole even when PLK4 remains as a ring after proteasome inhibition (Ohta et al., 2014), suggesting that STIL recruitment to a single site within the ring of Plk4 could even be the initial trigger for symmetry breaking in certain circumstances. In *Drosophila* S2 cells, the first observed break in symmetry is the recruitment of the STIL homologue, Ana2, to a single spot on the mother centriole (Dzhindzhev et al., 2017).

Is there a link between PLK4, Ana2 and the nuclear envelope? In various cell types, including *Drosophila* syncytial embryos, the centrosomes are tightly associated with the nuclear envelope via interactions between the microtubules they organise and nuclear-envelope-associated Dynein (Agircan et al., 2014; Bolhy et al., 2011; Raaijmakers et al., 2012; Robinson et al., 1999; Splinter et al., 2010). From our observations, we speculate that molecules associated with the nuclear envelope or concentrated within the local environment between centrosomes and the nuclear envelope may help determine the position of centriole duplication proteins in *Drosophila* syncytial embryos. These putative molecules may help stabilise Plk4 or recruit Ana2, or both. This could relate to the asymmetry in centrosomal microtubules, with differences in the ability of the microtubules connecting the centrosomes to the nuclear envelope and the microtubules extending out into the cytosol to concentrate PLK4 and Ana2. Alternatively, perhaps proteins associated with the nuclear envelope can transiently bind Plk4 or Ana2 and thus increase their local concentration in the region between the mother centriole and nuclear envelope. Ana2 directly interacts with a conserved member of the Dynein complex, Cut-up (Ctp), which is a form of Dynein Light Chain in *Drosophila* (Slevin et al., 2014; Wang et al., 2011). Although the precise function of the Ana2-Ctp interaction remains unclear, it appears to help mediate Ana2 tetramerisation (Slevin et al., 2014), and Ana2 tetramerisation is important for centriole assembly (Cottee et al., 2015). Thus, while Ctp does not appear to be essential for centriole duplication (Wang et al., 2011), any Ctp molecules released from the nuclear associated Dynein complexes would be ideally positioned to bind to Ana2 and promote daughter centriole assembly on the side of the mother centriole facing the nuclear envelope. These ideas are speculative and further work is needed to understand the molecular basis for the positional bias, as well as understanding its importance, if any. It will also be interesting to see whether positional bias occurs in other systems. Intriguingly, LRRCC1 has recently been shown to localise asymmetrically within the lumen of human centrioles with the position of procentriole assembly being non-random with respect to this asymmetric mark (Gaudin et al., 2021). Thus, although the molecular nature may vary, it’s possible that a non-random positional preference in daughter centriole assembly is an important conserved feature of centriole duplication.

## Acknowledgements

This work was supported by a BBSRC New Investigator Award (BB/P019188/1), a Wellcome Trust and Royal Society Sir Henry Dale Fellowship (105653/Z/14/Z) and an IdEx Université de Paris ANR-18-IDEX-0001 awarded to PTC. We thank Jordan Raff for fly lines and the use of his spinning disk microscope. We thank Corinne Tovey for critical reading of the manuscript. The work benefited from use of the imaging facility at the Stem Cell Institute, University of Cambridge and the imaging facility at the Institut Jacques Monod, Université de Paris. NHJC made the initial observation of positional preference by performing and analysing dual FRAP experiments with Spd-2-GFP / Sas-4-mCherry and Spd-2-GFP / RFP-Cnn, devised the formula to calculate angles, and performed and analysed the live Airyscan experiments. IB collected additional data for the dual FRAP experiments with Spd-2-GFP and Sas-4-mCherry and measured distances between centrioles in detached versus attached centrioles. PTC designed the study, performed all other experiments and analysis, and wrote the manuscript. The authors declare no financial or non-financial competing interests.

## Materials and methods

### Contact for Reagent and Resource Sharing

Further information and requests for resources and reagents should be directed to and will be fulfilled by the Lead Contact, Paul Conduit.

### Experimental Model and Subject Details

All fly strains were maintained at 18 or 25°C on Iberian fly food made from dry active yeast, agar, and organic pasta flour, supplemented with nipagin, propionic acid, pen/strep and food colouring.

### Methods

#### *Drosophila melanogaster* stocks

The following fluorescent alleles were used in this study: pUbq-Spd-2-GFP (Dix and Raff, 2007), eSas-4-mCherry (endogenous promoter) (Conduit et al., 2015a), pUbq-RFP-Cnn (Conduit et al., 2010), eSas-4-GFP (endogenous promoter) (Novak et al., 2014), eAsl-mCherry (endogenous promoter) (Conduit et al., 2015a), pUbq-GFP-Cnn-T1133A (this study). To make the pUbq-GFP-Cnn-T1133A allele, we used QuikChange (Agilent) to introduce the T1133A mutation into Cnn within a pDONR vector and used Gateway cloning (ThermoFisher) to transfer it into a pUbq-GFP vector containing a miniwhite marker. This construct was injected by BestGene in order to generate transgenic lines.

For performing FRAP experiments we used fly lines expressing either: two copies of pUbq-Spd-2-GFP and two copies eSas-4-mCherry in a *sas-4* null background (sas-4^I(3)2214^/Df(3R)BSC221); two copies of pUbq-Spd-2-GFP and one copy of RFP-Cnn in a cnn^f04547^/ cnn^HK21^ mutant background; two copies of eSas-4-GFP and two copies of eAsl-mCherry in a *sas-4* null background (sas-4^I(3)2214^/Df(3R)BSC221); or one copy of pUbq-GFP-Cnn-T1133A and two copies eSas-4-mCherry in a *sas-4* null background (sas-4^I(3)2214^/Df(3R)BSC221). For the live Airyscan imaging, we used flies expressing two copies of pUbq-Spd-2-GFP and two copies eSas-4-mCherry in a *sas-4* null background (sas-4^I(3)2214^/Df(3R)BSC221). For the fixed Airyscan imaging, we used flies expressing one copy of pUbq-GFP-Cnn-T1133A and two copies eAsl-mCherry in an *asl* null mutant background (asl^mecd^ (Blachon et al., 2008)).

#### Fixed and live cell imaging

For live dual FRAP experiments, 0.5μm thick confocal sections were collected from living syncytial embryos in nuclear cycle 11 or 12 at ∼21°C on either a Perkin Elmer ERS Spinning Disk confocal system mounted on a Zeiss Axiovert microscope using a 63X/1.4NA Oil objective, or an Andor Revolution Spinning Disk confocal system mounted on a Nikon Ti inverted microscope coupled to an Andor iXon camera using a Plan-Apochromat 60X/1.4NA Oil objective. Focused 488nm and 561nm lasers were used to photobleach the GFP and mCherry/RFP signals, respectively. For live Airyscan imaging, 0.2 μm thick sections were collected from living embryos in nuclear cycle 12 or 13 on an inverted Zeiss 880 microscope fitted with an Airyscan detector at 21°C and a Plan-Apochromat 63×/1.4NA oil lens using 488-nm argon and 561-nm diode lasers. Images were collected approximately every 1 min with a zoom value of 23.3 pixels/μm. Focus was readjusted between the 1-min intervals. Images were Airy-processed in 3D with a strength value of “auto” (∼6) or 6.5. For fixed Airyscan imaging, 0.2 μm thick sections were collected from methanol fixed embryos in nuclear cycle 11 or 12 on an inverted Zeiss LSM980 microscope fitted with an Airyscan2 detector at 21°C and a Plan-Apochromat 63×/1.4NA oil lens using 488-nm argon and 561-nm diode lasers. When measuring centriole positions, images from the different colour channels were registered with alignment parameters obtained from calibration measurements with 0.2 μm diameter TetraSpeck beads (Life Technologies). The centroids of each fluorescent signal were calculated in ImageJ using the “centre of mass” analysis tool. The number of pixels for the images was first increased such that each real pixel was made of 5×5 sub-pixels. This increases the location accuracy for the centroid of the fluorescence signal.

#### Quantification and Statistical Analysis

Data was processed in Microsoft Excel. Graph production was performed using either Microsoft Excel (rose plots) or GraphPad Prism (all other graphs) and statistical analysis was performed using GraphPad Prism. N numbers and statistical tests used for each experiment are indicated within the main text or Figure Legends. The following Normality tests were carried out in Prism to analyse the frequency distributions of angles: Anderson-Darling test, D’Agostino & Pearson test, Shapiro-Wilk test, Kolmogorov-Smirnov test.

